# Brain correlates of action word memory

**DOI:** 10.1101/412676

**Authors:** Zubaida Shebani, Francesca Carota, Olaf Hauk, James B. Rowe, Lawrence W. Barsalou, Rosario Tomasello, Friedemann Pulvermüller

## Abstract

When understanding language semantically related to actions, the motor cortex is active and may be sensitive to semantic information, for example about the body-part-relationship of displayed action-related words. Conversely, movements of the hands or feet can impair memory performance for arm- and leg-related action words respectively, suggesting that the role of motor systems extends to verbal working memory. Here, we studied brain correlates of verbal memory load for action-related words using event-related fMRI during the encoding and memory maintenance of word lists. Seventeen participants saw either four identical or four different words from the same category, semantically related to actions typically performed either with the arms or with the legs. After a variable delay of 4-14 seconds, they performed a nonmatching-to-sample task. Hemodynamic activity related to the information load of words at presentation was most prominent in left temporo-occipital and bilateral posterior-parietal areas. In contrast, larger demand on verbal memory maintenance produced greater activation in left premotor and supplementary motor cortex, along with posterior-parietal areas, indicating that verbal memory circuits for action-related words include the cortical action system. Somatotopic memory load effects of arm- and leg-related words were not present at the typical precentral loci where earlier studies had found such word-category differences in reading tasks, although traces of somatotopic semantic mappings were observed at more anterior cortical regions. These results support a neurocomputational model of distributed action-perception circuits (APCs), according to which language understanding is manifest as full ignition of APCs, whereas working memory is realized as reverberant activity gradually receding to multimodal prefrontal and lateral temporal areas.

## 1. Introduction

When reading and listening to action words, humans automatically think of the respective action. This recognition of action words is accompanied by the instantaneous neurophysiological activation of motor systems (Hauk and Pulvermüller, 2004; Barsalou, 2008; Pulvermüller and Fadiga, 2010; Willems and Casasanto, 2011; Glenberg and Gallese, 2012; Kiefer and Pulvermüller, 2012; Klepp et al., 2014). The reverse functional link between action and language systems is shown by behavioral and TMS studies in which motor system activity modulates the processing of action words (Pulvermüller et al., 2005; Gerfo et al., 2008; Willems et al., 2011; Repetto et al., 2013; Gianelli and Dalla Volta, 2015; Vukovik et al., 2017; Shebani and Pulvermüller, 2018). For example, stimulating the motor cortex using TMS modulates the recognition of semantically-specific types of action words (Pulvermüller et al., 2005; Willems et al., 2011) and motor movement can interfere with or facilitate action word processing and memory (Glenberg and Kaschak, 2002; Glenberg et al., 2008; Shebani and Pulvermüller, 2013; 2018) just as the processing of action related words and sentences can interfere with or assist motor movement (Boulenger et al., 2006; de Vega et al., 2013). Additionally, dysfunction of motor systems found with focal cortical damage or more widespread progredient disease impairs the processing of action words and concepts (Bak et al., 2001; Cotelli et al., 2006; Boulenger et al., 2008; Bak and Chandran, 2012; Kemmerer et al., 2012; Fernandino et al., 2013).

Together, these results demonstrate a causal meaning-dependent influence of motor systems on symbol processing and lead to the hypothesis that a network of interacting areas contributes to both action-semantics and symbolic-linguistic processes for the perception and comprehension of action-related words; the contribution of motor areas has been proposed to be crucial because it provides the necessary semantic grounding of the linguistic symbols in bodily action (Barsalou, 2008; Pulvermüller, 2013). The same network of interacting areas involved in action word comprehension has also been suggested to be relevant and critical for working memory processes.

Here we ask whether memory maintenance for different kinds of action related words would draw on the same areas also involved in language comprehension and in the motor actions these words are used to speak about. Working memory refers to the retention and processing of information that is just experienced but no longer available in the external environment, or to information retrieved from long-term memory (Fuster, 1995; D’Esposito, 2007). Over the past 30 years, several cognitive models of working memory have been proposed (Baddeley, 1992; Cowan, 1998). Baddeley’s highly influential model of working memory (e.g., Baddeley, 2003) includes a ‘central executive’ to control attention and to manage information in verbal and visuospatial buffers. Internal representations held in working memory can be actively maintained through rehearsal strategies mediated by sub-vocal articulation. Verbal working memory engages a network of brain regions thought to be involved in articulatory and auditory phonological processing, including inferior frontal (Broca’s area) and superior temporal cortex (Wernicke’s area) along with parietal cortex (Paulesu et al., 1993; Schumacher et al., 1996; Buchsbaum and D’Esposita, 2008). A key region associated with verbal memory tasks is the dorsolateral prefrontal cortex (PFC), which is sometimes presented as the locus of the ‘frontal executive’ functions (Baddeley, 2003). Different areas of PFC support different memory sub-functions (memory maintenance to BA 9 and memory-response selection to BA 46, Rowe et al., 2000). Convergent input to this region from sensory, motor areas and association cortex may explain why this region plays such a prominent role in memory processes (Fuster, 1997; 2015). In this sense, working memory may not just be the product of the PFC but of its interactions with posterior cortical areas (Fuster, 1995; Petrides, 2000; D’Esposito, 2007; Rowe et al., 2007; Fuster, 2009).

The neurobiological mechanisms of working memory have been elucidated by intracortical recordings. From this research, it emerged that neurons in prefrontal cortex and in multimodal parietal and temporal regions are most likely to include memory cells indicating specific content relevant in working memory tasks. Interestingly, parallel cell dynamics and memory-content specificities were found in frontal and temporal systems and temporary lesions in one of these systems were observed to entail functional changes in the other (for review, see Fuster, 1995; 2003; 2009). This body of evidence enforces the position that it is not areas that are responsible for working memory but neuronal ensembles, called action perception circuits (APCs), whose strongly interlinked neuron members are distributed across several areas and maintain their reverberant activity for some time. Mathematical models of brain function support this position (Verduzco-Flores et al., 2009). Neurocomputational modelling of word learning show the emergence of action perception circuits distributed across language areas, including the inferior-frontal and superior-temporal language areas of Broca and Wernicke (Garagnani et al., 2008; Pulvermuller, 2018). When word meaning is grounded in action and perception, these linguistic circuits link up with additional motor and sensory circuits (Garagnani and Pulvermüller, 2016; Tomasello et al., 2017). This leads to an extension of symbolic networks into sensorimotor areas, including motor cortex. Note that, although the “grounding” of semantic knowledge requires participation of sensorimotor areas, the resultant APCs are not restricted to such modality-specific areas. Because activity spreading between sensory and motor fields requires passing-though other areas, in particular multimodal ones with high degree of connectivity (so-called ‘connector hubs’), the APCs span across relevant modality-preferential and -general regions (Garagnani et al., 2008).

Neuronal circuit models explain why language and symbol processing activate classic language areas along with multimodal prefrontal and temporal areas, and even, depending on semantic word type, additional category-preferential areas such as the motor cortex. However, previous model simulations make an additional important prediction on how APC dynamics change over time. After initial full activation (‘ignition’ Braitenberg, 1978; Palm et al., 2014; Moutard et al., 2015) involving the entire APC, activity within the circuit decreases due to neuronal inhibition and fatigue; only neurons in those network parts most strongly interlinked with other circuit members are able to maintain activity over several seconds, and therefore contribute to working memory. These neurons are primarily in the strongly connected ‘connector hubs’. Therefore, after ignition, activity retreats from the modality-preferential areas relevant for grounding to multimodal connector hubs. In the frontal cortex, as shown in Figure 1, this would result in an anterior shift from motor cortex to adjacent frontal and prefrontal cortex (Pulvermüller and Garagnani, 2012; Tomasello et al., 2017; Pulvermuller, 2018). A strong version of an embodied perspective on semantic meaning may put that, similar to symbolic understanding, the memory maintenance of action words draws primarily on motor systems. In addition to these three competing predictions, we explored whether similar body part specific motor regions as previously shown during word perception and understanding are active during the retention of action words in working memory.

**Figure 1.**
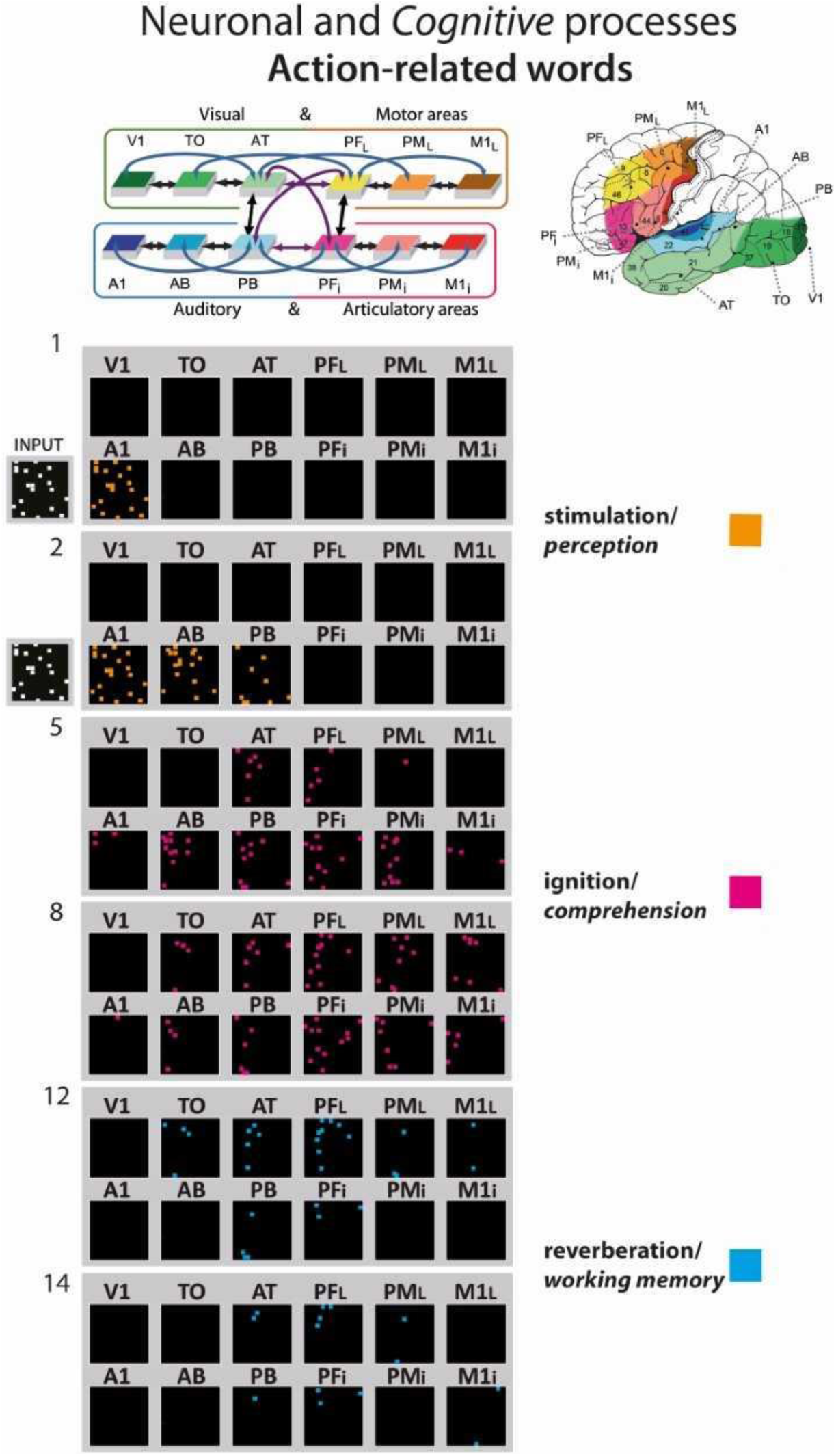
Simulated action word processing in a biologically constrained spiking network model of the fronto-temporo-occipital lobes. After the network underwent action word learning by interlinking acoustic, articulatory and action-semantic information, the action-word-related circuit was re-activated by auditory stimulation to areas A1. The re-activation process comes in different consecutive neuronal and cognitive phases, the stimulation phase, which corresponds to word perception (orange pixel), the full activation or ‘ignition’ phase, the correlate of word comprehension (magenta pixel), and the reverberant maintenance of activity, which underpins verbal working memory (blue pixels). Please note the relatively prominent role of prefrontal cortex in the reverberation and working memory phase, which motivates the prediction of an anterior frontal activity shift. At the top right, the 12 brain areas modelled are shown. The top left box-and-arrow diagram shows the structure of the network; box colours and positions indicate correspondence to brain area and arrows between area connectivity. Sets of 12 black squares in the main diagram below represent activation of the same 12 areas at a given simulation time step. Simulation time steps are indicated on the left. Each coloured dot represents one active (spiking) model neuron at a given time step. Data from Tomasello et al. (2017; submitted).

In the present study, we used arm and leg related action words to examine brain correlates of verbal working memory. A low load condition with four repetitions of the same to-be-memorised word was compared with a high load condition with four different words that are semantically closely related. Hemodynamic responses were obtained when subjects read and encoded word stimuli and subsequently when they maintained them in their working memory. Long and variable memory delays reduced the correlations between these haemodynamic correlates of neural activation. We expected that the activation of 1) Broca’s area, 2) motor regions or 3) frontal areas anterior to the motor regions previously found active during word comprehension would characterize the high-load memory maintenance period (as compared with the low load condition and the memory encoding phase). Furthermore, we asked whether semantic differences between word types, namely their respective relationship to upper and lower extremities, might lead to category-specific activations reminiscent of the semantic somatotopy found in word reading or recognition experiments.

## 2. Materials and Methods

### 2.1 Participants

Nineteen monolingual, native English speakers participated in the study. All were right handed with an average laterality quotient (Oldfield, 1971) of 74.9% (s.d. = 22.6). All participants gave their written, informed consent and were reimbursed for their time. One subject was discarded prior to statistical analysis of fMRI data due to excessive movement during the acquisitions (more than 10mm). A further subject was discarded due to poor performance on the behavioural task (50% errors). Therefore, data from 17 subjects (9 male; aged 21-35, mean 25.5, SD 3.8) are reported below. All participants had normal or corrected-to-normal vision and confirmed that they were without psychiatric or neurological illnesses and did not use any medication or drugs. Ethics permission for the study was obtained from the Cambridge Psychology Research Ethics Committee.

### 2.2 Stimuli

Lexical stimuli for the task consisted of 80 action words, 40 semantically related to the arm (e.g., “pick”, “grasp”) and 40 to the leg (“walk”, “kick”). They were matched for a number of psycholinguistic variables (see Table 1) including word, lemma, bigram, and trigram frequencies, and their number of letters and phonemes. Lexical stimuli were also matched for grammatical ambiguity, and for ratings of valence, arousal, imageability, visual relatedness, body relatedness and general action relatedness as revealed by previous semantic ratings (Hauk and Pulvermüller, 2004). In addition to the original set of 80 arm/leg words, 10 action words (5 arm/5 leg) were used as probes in non-match trials. Two different pseudo-randomised stimulus sequences with the same repetition structure for arm and leg words were used and alternated between subjects.

**Table 1.** Means and standard errors of psycholinguistic and semantic properties for arm and leg words. Differences between arm and leg words were n.s. at p < 0.05.

| Variable | Arm words |  | Leg words |  |
| --- | --- | --- | --- | --- |
|  | Mean | SE | Mean | SE |
| Number of phonemes | 3.73 | (0.12) | 3.90 | (0.14) |
| Number of letters | 4.45 | (0.14) | 4.57 | (0.13) |
| Grammatical ambiguity | 1.93 | (0.04) | 1.95 | (0.03) |
| Word frequency | 219.8 | (47.0) | 232.8 | (48.2) |
| Lemma frequency | 520.2 | (82.1) | 540.5 | (89.9) |
| Bigram frequency | 30196 | (2506) | 34859 | (2726) |
| Trigram frequency | 3250 | (386.4) | 3076 | (317.2) |
| Valence | 3.65 | (0.14) | 3.96 | (0.14) |
| Arousal | 3.04 | (0.14) | 3.12 | (0.16) |
| Imageability | 4.60 | (0.12) | 4.53 | (0.14) |
| Visual relatedness | 4.40 | (0.16) | 4.14 | (0.16) |
| Body relatedness | 3.71 | (0.16) | 3.74 | (0.14) |
| Action relatedness | 5.06 | (0.14) | 5.11 | (0.17) |

### 2.3 Procedure

During MRI scanning, subjects performed a delayed non-match-to-sample task consisting of four blocks. In each trial, subjects were presented with either four arm- or leg-related action words and, after a delay of 4-14 seconds – the memory period – they were shown one more word – the probe. The task required subjects to press a button with their left index finger whenever the probe word was *different* from all of the four previous words in that trial; subjects had to rest and avoid any movements when probe stimuli matched one of the four sample stimuli. Sample words were presented serially for 200 ms each, with a stimulus onset asynchrony (SOA) of 500 ms. Subjects were instructed to keep all of the four sample words in memory during the subsequent delay period and to respond to the probe as fast and as accurately as possible. The length of the delay varied randomly between 4 and 14 s. After the delay, the probe word, which was from the same action word category as the sample stimuli of a given trial, was shown for 1 s. In order to minimise movements in the scanner, subjects were instructed to respond by button press in non-match trials only, which constituted 20% of all trials. Left hand responses were required to minimise motor-related activation in the left language-dominant hemisphere, where relevant language related activations were expected. Subjects had up to 4 s to respond to the probe in each trial. Note that probe words in non-match trials were different arm/leg-related action words than those in the original set of 80 action words and never presented in the task as sample stimuli. A variable-length inter-trial interval (ITI) (8-12 s, counterbalanced) separated all trials. The central fixation cross present during the inter-stimulus interval and the memory period was olive in colour then changed to grey during the ITI. The fixation cross changed from grey back to olive for 1 s at the end of the ITI to alert subjects to the beginning of the upcoming trial.

Memory load was varied between trials. In the high load condition, four different action words, which were also very close in meaning, were presented, whereas in the low load condition a single action word was shown four times. Each block consisted of 40 trials, 20 with arm- and 20 with leg-related action words, each group again subdivided into 10 high and 10 low load trials. Trials were pseudo-randomised within each block so that not more than two trials of the same action word category (arm/leg) appeared consecutively.

Before scanning, subjects viewed task instructions and performed a practice version of the task. Responses and reaction times were recorded using an MRI compatible button box. The task was designed and presented and behavioural data was recorded using E-Prime 1.1 (Psychology Software Tools Inc., Sharpsburg, BA, USA).

### 2.4 fMRI Data Acquisition

Participants were scanned on a Siemens (Erlangen, Germany) TIM Trio 3T machine at the MRC Cognition and Brain Sciences Unit (MRC-CBSU), Cambridge, UK. The nonmatching-to-sample task was performed during 4 separate sessions of echoplanar imaging (EPI) with 460 volumes acquired in each session (including 12 s of initial dummy scans to allow steady state magnetisation). Acquisition parameters used were as follows: TR = 2.02 s; TE = 30 ms; flip angle = 78°. Functional images consisted of 32 interleaved slices covering the whole brain (slice thickness 3mm; matrix size 64 x 64; interslice gap 25%; in-plane resolution 3 x 3 mm; see http://imaging.mrc-cbu.cam.ac.uk/imaging/ImagingSequences). Stimuli were back-projected onto a screen using a Christie video projector with a 60-Hz refresh rate, and viewed using a mirror mounted on the head-coil. Soft padding minimized head movement during the scanning session.

### 2.5 fMRI Data Analysis

Imaging data were processed and analysed using the SPM5 (Wellcome Department of Imaging Neuroscience, London, UK; http://www.fil.ion.ucl.ac.uk/spm/). Images were first corrected for slice timing, and then realigned to the first image using sinc interpolation. Any non-brain parts were removed from the T1-weighted structural images using a surface model approach (Smith, 2002). The EPI images were coregistered to these structural T1-images using a mutual information coregistration procedure (Maes et al., 1997). The structural MRI was then normalized to the 152-subject T1 template of the Montreal Neurological Institute (MNI). The resulting transformation parameters were applied to the coregistered EPI images. During spatial normalization, images were re-sampled with a spatial resolution of 2 × 2 × 2 mm^3^ and spatially smoothed with a 10 mm full-width half-maximum Gaussian kernel. Preprocessing was automated using in-house software (http://imaging.mrc-cbu.cam.ac.uk/imaging/AutomaticAnalysisManual).

Individual subject activations were analysed using a general linear model approach (Friston et al., 1998). A high-pass filter was used to remove low-frequency noise in the signal (cutoff period 128 s). The data for each subject were modelled using a boxcar design convolved with the canonical haemodynamic response function. Events of interest and time points modelled were as follows: encoding (0-2 s), memory period (2-11 s +/-5 s, adjusted to the length of individual memory periods) and two probe events at the end of the memory period, one for trials requiring a response and one for non-response trials. This generated a time-course of predicted neural activity for each event type allowing us to estimate changes in haemodynamic signal for arm/leg word stimuli in the high and low load memory conditions. Four stimulus events (hi/lo memory load condition, arm/leg words) were distinguished in the encoding and memory maintenance intervals respectively; additional response and non-response events were coded for the final retrieval interval. Contrasts were run to estimate signal changes associated with these events at each voxel and the resulting maps from each subject were entered into a second level (group) analysis treating subjects as a random variable. Brain activations are displayed after controlling for false discovery rate (FDR) at 0.05 for multiple comparisons. Stereotaxic coordinates for voxels with maximal z values within activation clusters are reported in MNI standard space. Anatomical labels of nearest cortical grey matter for peak coordinates were obtained from the MRIcron software (http://www.sph.sc.edu/comd/rorden/mricro.html), based on the anatomical parcellation of the MNI brain published by (Tzourio-Mazoyer et al., 2002).

### 2.6 ROI analyses

In addition to the whole brain analysis, activity in regions of interest (ROI) was examined. One such analysis focused on activation differences between the initial memory encoding interval and the subsequent memory maintenance epoch. Further analyses were performed to compare memory load effects for arm and leg related action words. For data driven ROI definition, clusters activated due to memory load (encoding and maintenance periods together) using a whole-brain corrected significance criterion were used. Each absolute activation maximum (that is, the voxel with the highest t-value in its respective significant cluster) was defined as the centre of an ROI with radius 10 mm. The MarsBar software utility (http://marsbar.sourceforge.net/) was used to average parameter estimates over voxels and to estimate signal changes in these regions for each time interval (encoding, maintenance), word type (arm, leg) and subject. These data were then submitted to a repeated measures analysis of variance (ANOVA). F-Tests, Bonferroni-corrected for multiple comparisons, were used as planned comparison tests.

In order to examine whether working memory produces activation in regions anterior to those found for action word perception, four additional ROIs were selected based on local activation maxima in frontocentral sensorimotor cortex during encoding and memory. Two lateral and two dorsal precentral ROIs were contrasted. These were 1.3-3cm anterior-lateral to regions where previous studies had found word-category differences in brain activation in reading and listening tasks (see Results, cf. Carota et al., 2012). Activation in these regions was compared between word categories using an additional ANOVA.

## 3. Results

### 3.1 Behaviour

High accuracy rates (mean = 97.7%, standard error, SE = 0.3%) and d’ values (mean = 3.9, SE = 0.12) confirmed good performance in all subjects.

### 3.2 Whole brain analysis

The memory load contrast (high load vs. low load) showed significant activation (*p* < 0.05, FDR corrected) in a range of areas (Table 2, Figure 2, top and bottom right panels). One activated cluster appeared in left precentral gyrus (BA6), also extending to adjacent motor and prefrontal cortex (caudal BA8, 9). This cluster stretched from dorsolateral sites down into the posterior, premotor part of Broca’s area (BA44/45). A second cluster was in left dorsal supplementary motor area, SMA (BA6), and a third extended from left inferior-occipital cortex into inferior-temporal and superior-temporal sites. A fourth cluster in the left hemisphere included the intraparietal sulcus and adjacent temporal areas (BA 40, 7). Right hemispheric activation indicative of memory load was seen in precentral/prefrontal and parietal areas homotopic to the left-hemispheric ones. In addition, right inferior temporo-occipital cortex and possibly adjacent cerebellum showed general memory load effects.

**Figure 2.**
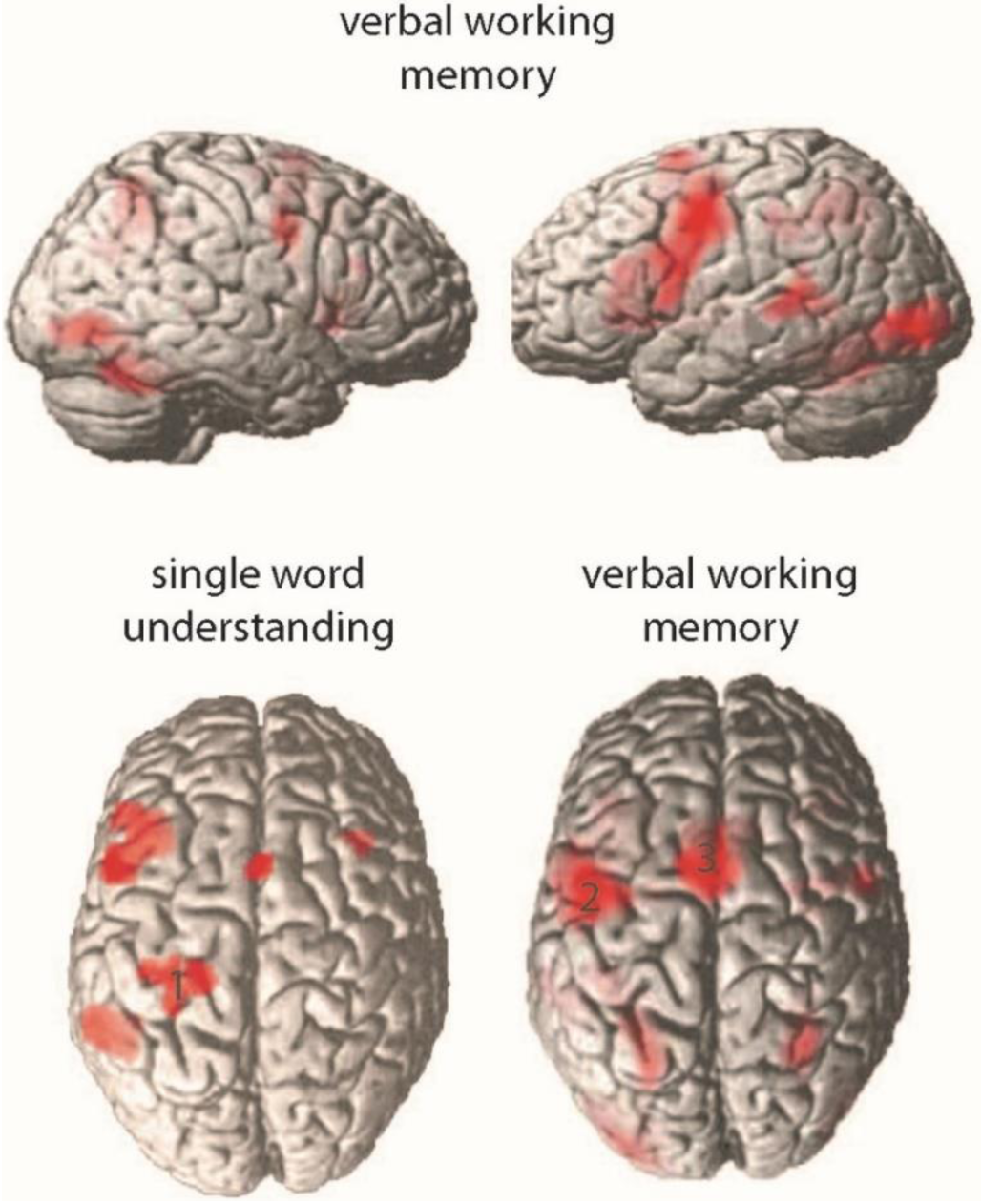
Top panels: Hemodynamic correlates of verbal memory load in the delayed nonmatching-to-sample task. Trials with high memory load are compared against a baseline of low memory load, while keeping constant both task and amount of stimulation. Both encoding and memory maintenance intervals are collapsed into this analysis. All clusters are significant at an FDR-corrected threshold p < 0.05. Bottom panels: Dorsal views of BOLD activation during passive reading of action words (against a baseline of looking at matched meaningless symbol strings; bottom left) and of the memory load contrast (as in top panels). Note the central position of the activation focus labelled ‘1’ in sensorimotor cortex in the former and the more anterior foci labeled ‘2’ and ‘3’ in lateral and dorsomedial frontal cortex in the latter. Note also that the anterior left inferior prefrontal activation focus in the former (bottom left) is largely due to the face related words included in the study (Hauk et al., 2004), which were not used in the present study.

**Table 2.**
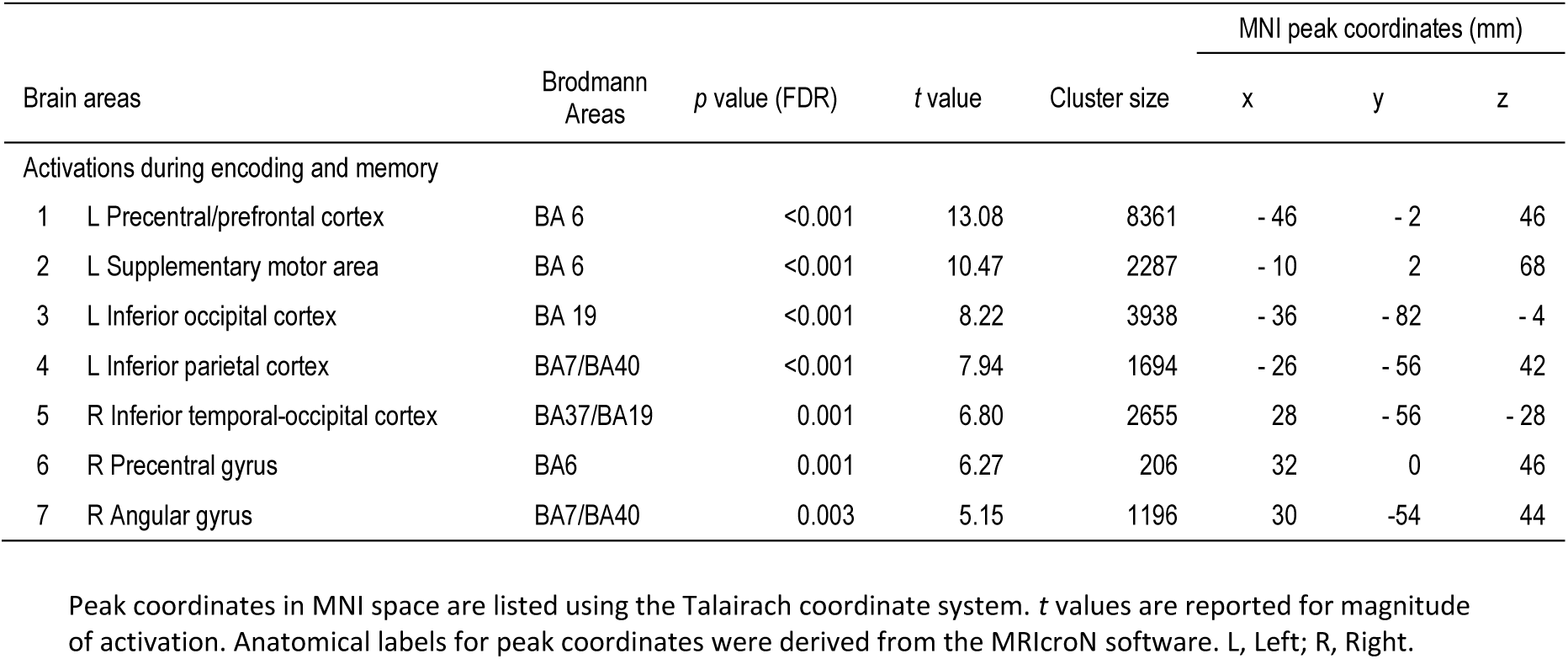
Significant areas of activation during encoding and memory (threshold at FDR 0.05).

To assess the hypothesis of an anterior shift of frontal activity during the present working memory task as compared with a passive reading and understanding paradigm, we contrast the results reported by Hauk et al. (2004, their figure 1C, left panel), with the whole brain analysis of the present study (Figure 2, bottom panels). It can be seen that, instead of a pronounced activation focus extending across the central sulcus (labeled by the number ‘1’), which was present in the word comprehension study, the current high vs. low memory load contrast showed two significant foci, one encompassing lateral premotor and posterior prefrontal cortex and one including dorsomedial SMA (which are labeled ‘2’ and ‘3’). This result provides clear evidence for an anterior shift in frontal activity in working verbal memory for action words.

Whole brain analyses performed separately on the memory load contrasts obtained for the initial stimulus encoding interval and on those for the subsequent period of active memory maintenance indicated differences between these time periods. As Figure 3 shows, stimulus encoding (in red) activated a range of areas, including temporo-occipital, superior-temporal and intra-parietal sites. Active memory maintenance (in green) produced activation in inferior-parietal and superior-temporal regions. A list of significant areas of activation for each time interval is presented in Table 3.

**Figure 3.**
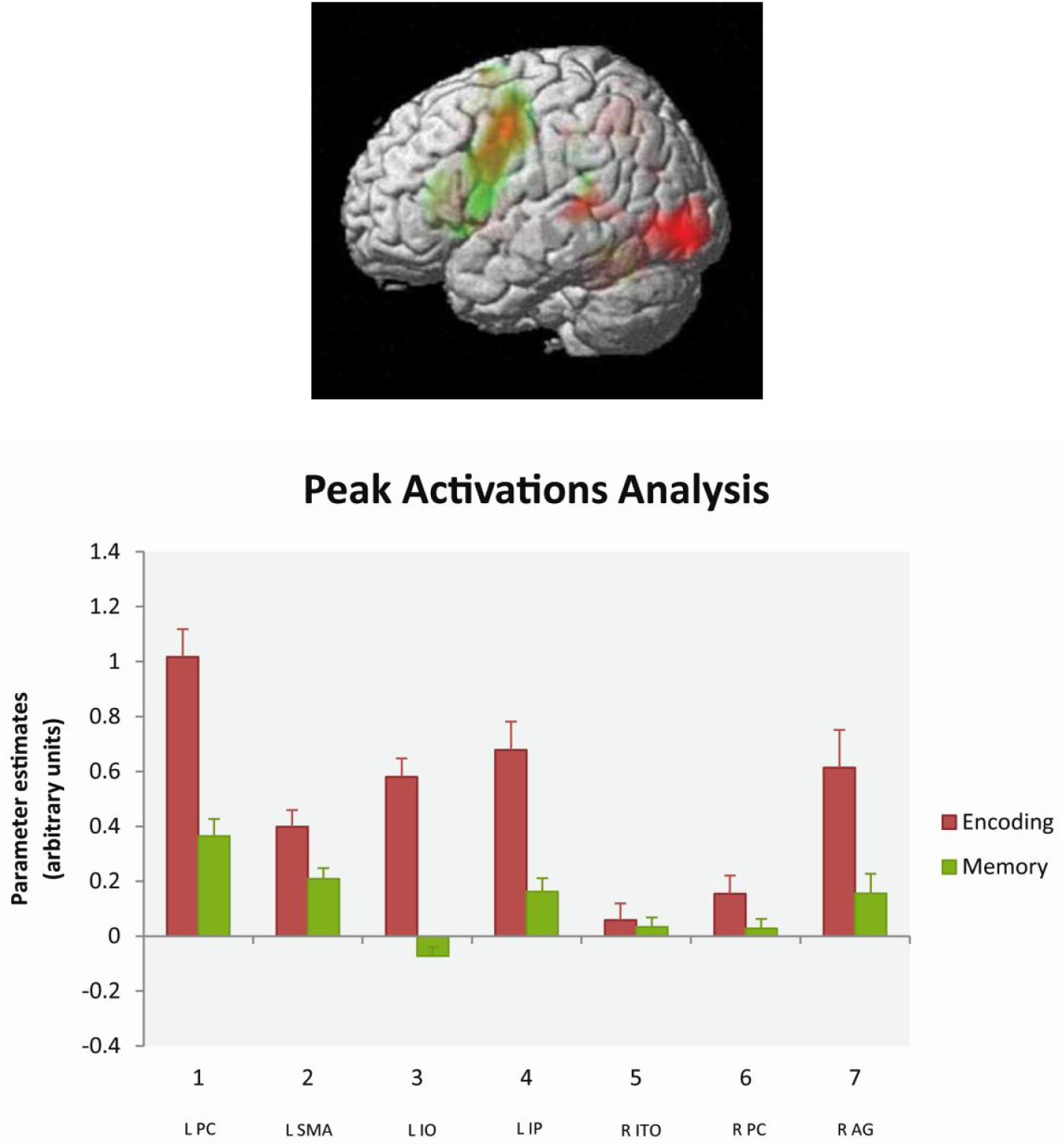
Memory load effects during the encoding interval (in red) contrasted with that during memory maintenance (in green; FDR p < 0.05). Note the pronounced overlapping activation in left dorsolateral premotor/prefrontal cortex and in the supplementary motor area. L PC, left precentral/prefrontal cortex; L SMA, left supplementary motor area; L IO, left inferior occipital cortex; L IP, left inferior parietal cortex; R ITO, right inferior temporal-occipital cortex; R PC, right precentral gyrus; R AG, right angular gyrus.

**Table 3.** Significant areas of activation during the encoding, memory and retrieval time intervals in the memory load contrast (threshold at FDR 0.05).

| Brain areas | *p* value (FDR) | *t* value | Cluster size | MNI peak coordinates (mm) |  |  |
| --- | --- | --- | --- | --- | --- | --- |
|  |  |  |  | x | y | z |
| Activations during encoding |  |  |  |  |  |  |
| L Precentral gyrus | <0.001 | 11.16 | 4887 | - 46 | - 2 | 46 |
| L Inferior occipital cortex | <0.001 | 9.87 | 3577 | - 30 | - 88 | - 8 |
| R Inferior occipital cortex | <0.001 | 7.76 | 2645 | 48 | - 76 | - 6 |
| L Supplementary motor area | <0.001 | 7.69 | 1778 | - 8 | 0 | 68 |
| L Superior parietal cortex | <0.001 | 7.28 | 1189 | - 24 | - 56 | 44 |
| R Precentral gyrus | 0.002 | 5.81 | 538 | 56 | -2 | 44 |
| R Mid occipital | 0.003 | 5.25 | 731 | 34 | - 66 | 26 |
| R Putamen | 0.007 | 4.56 | 973 | 22 | 16 | 0 |
| R Calcarine | 0.022 | 3.53 | 36 | 16 | - 70 | 10 |
| R Mid frontal gyrus | 0.026 | 3.42 | 32 | 40 | 28 | 22 |
| Activations during memory maintenance |  |  |  |  |  |  |
| L Inferior frontal gyrus (operculum) | 0.014 | 7.95 | 8149 | - 62 | 8 | 8 |
| R Cerebellum | 0.016 | 6.55 | 1626 | 24 | - 60 | - 24 |
| L Supplementary motor area | 0.018 | 5.61 | 1028 | - 8 | 2 | 56 |
| L Superior temporal cortex | 0.024 | 4.62 | 126 | - 52 | - 42 | 18 |
| L Inferior temporal cortex | 0.027 | 4.24 | 201 | - 42 | - 46 | - 12 |
| R Caudate nucleus | 0.028 | 4.08 | 57 | 18 | 28 | 10 |
| R Insula | 0.030 | 3.89 | 201 | 34 | 18 | 8 |
| R Mid frontal | 0.031 | 3.76 | 39 | 32 | 40 | 30 |
| L Mid occipital cortex | 0.034 | 3.59 | 105 | - 24 | - 58 | 42 |
| Activations during retrieval |  |  |  |  |  |  |
| R Superior Frontal gyrus | 0.016 | 7.54 | 365 | 4 | 30 | 46 |
| L Precentral gyrus | 0.023 | 5.51 | 37 | - 54 | 12 | 32 |

Peak coordinates in MNI space are listed using the Talairach coordinate system. *t* values are reported for magnitude of activation. Only cluster sizes > 30 are presented. Anatomical labels for peak coordinates were derived from the MRICroN software and SPM 8. L, Left; R, Right.

### 3.3 ROI analyses

Somatotopic differences between memory load effects during encoding and active memory maintenance intervals was investigated further using a data-driven analysis of regions of interest (ROIs), which were placed around the peak activation voxels of all FDR corrected clusters of the general high-vs.-low-load contrast (Figure 2, Table 2). Average activation values obtained for each of these ROIs in each time interval (encoding vs. maintenance) and word type (arm-vs. leg-related) were submitted to an analysis of variance, ANOVA (with factors ROI, time interval and word type), which revealed a significant interaction between the factors ROI and Interval (F (6,96) = 14,21, p < 0.00001). Significant differences between time intervals were confirmed by planned comparison F-tests in left prefrontal/premotor, parietal and temporo-occipital along with right parietal ROIs (Bonferroni-adjusted significance threshold: p < 0.014). These data-driven ROIs showed relatively stronger activation during memory encoding. During the memory maintenance interval, activation was primarily observed in premotor and SMA regions (Figure 3).

The data-driven ROI analysis did not provide evidence for brain activation differences between word types. However, when rendering memory-load effects for arm and leg words separately at an uncorrected threshold of p < 0.001 (Figure 4), category differences were observed about 1-3cm anteriorly to where action word related differences had been reported previously (cf., Hauk, 2004; Kemmerer and Gonzalez-Castillo, 2010; Carota et al., 2012). Arm words (in red) activated inferior and lateral prefrontal and precentral areas, whereas leg words (in blue) activated additional more dorsal regions. Note that these tendencies towards differences were present not at the loci where strongest memory-load related activation was seen, but slightly anterior-lateral to these sites instead. In the second ROI analysis, defined around local activation maxima of the general memory load contrast (high-load-vs.-low-load) in which two lateral ROIs (-46 -2 44, -48 - 4 50) were contrasted with two dorsal precentral ROIs (-4 2 58, -34 0 60), the ANOVA with the design ROI × Word Category on the parameter estimates averaged over the voxels in each pair of the lateral and dorsal ROIs revealed a significant interaction of ROI and Word Category (F (1, 16) = 9.79, p= 0.0065) due to stronger leg-word than arm-word memory activation in the dorsal regions (t (1, 16) = 1.85, p < 0.04, one-tailed), but no significant differences at lateral sites.

**Figure 4.**
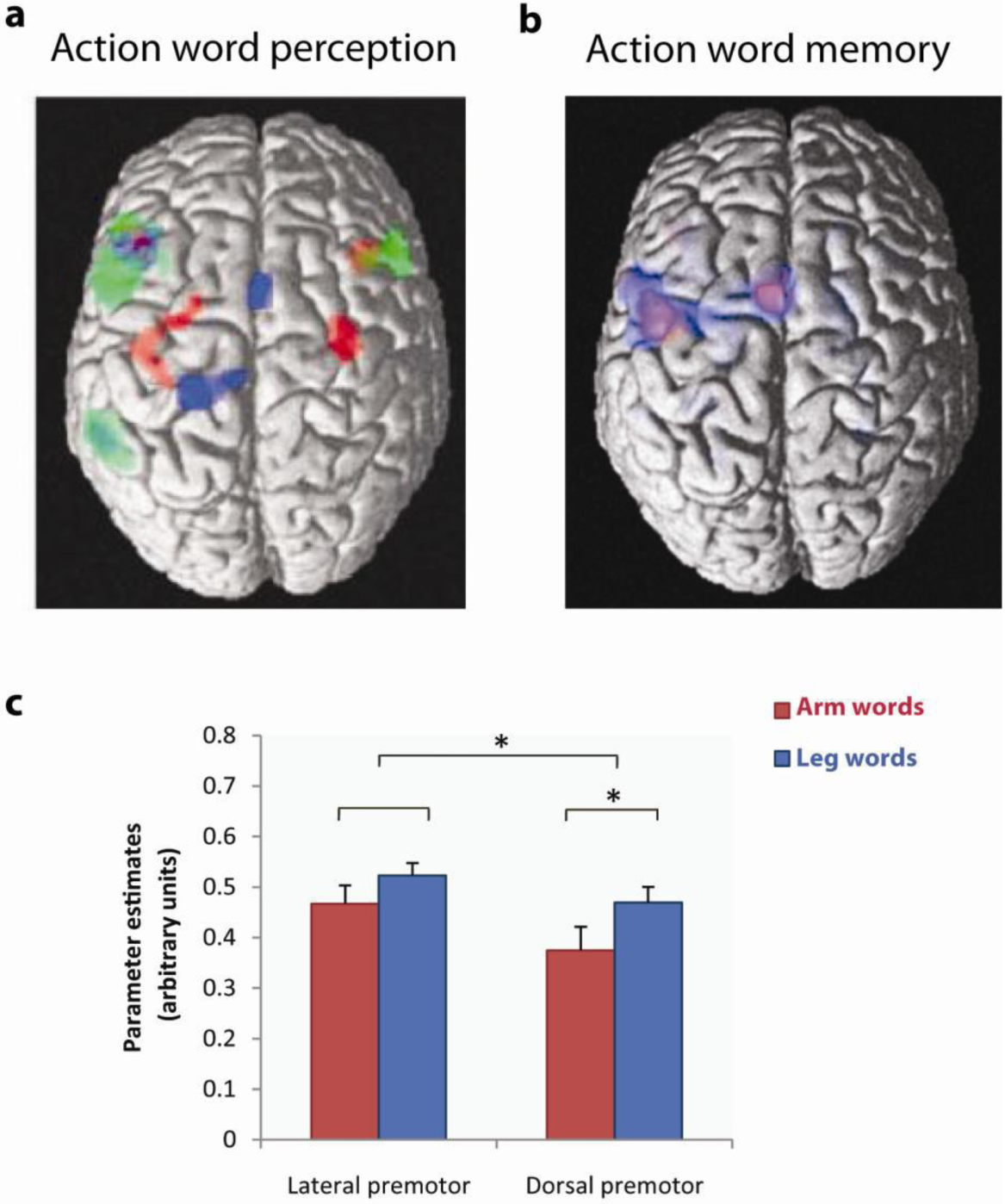
**(a, b)** Comparison between dorsal views of word category effects seen in the present working memory study and in an earlier study of word reading using a similar set of arm- and leg-related words (Hauk et al., 2004). The previous study’s results are displayed on the left, with activity to face-related words in green, that to arm-related words in red, and that to leg words in blue. The brain diagram on the right presents results on memory load effects from the present investigation (p < 0.001 uncorrected) with arm word-memory load highlighted in red and memory load for leg words in blue. Note the anterior shift of category-specific activation in verbal working memory relative to reading. **(c)** Significant interaction of ROI and Word Category in the present study showing stronger activation for leg-word than arm-word memory in dorsal premotor regions.

## 4. Discussion

The aim of the present study was to examine the brain correlates of verbal working memory for action-related words. More specifically, we investigated whether the same motor regions previously shown to be active during action word perception and understanding would remain active during memory maintenance, as predicted by the embodied perspective, or the main areas for verbal working memory as predicted by the Baddeley model with its emphasis on Broca’s region. As a third possibility, we considered the frontal memory shift hypothesis of current neurobiologically founded action perception theory, according to which active memory maintenance draws on multimodal connector hub areas and thus, in the case of action-related words, upon areas anterior to the motor regions typically found active during comprehension.

Our results provide unambiguous evidence for the anterior shift hypothesis and thus for the action perception model. As an additional feature, we asked whether semantic differences between words related to actions typically performed with different parts of the body might lead to category-specific activations resembling the semantic somatotopy found in word reading or recognition experiments.

Our results show memory load effects for arm- and leg-related action words in partially overlapping areas, with only weak evidence of category-specificity, anterior to the precentral sites previously associated with differences between semantic action word types. Please note once again that these results found in a memory task contrast with the previously reported pattern of activation during passive reading and listening to words. In these passive perceptual tasks, premotor and motor cortex showed meaning-related activation of arm and leg motor representations to arm- and leg-related words in a semantically somatotopic manner (Pulvermüller et al., 2001; Hauk and Pulvermüller, 2004; Shtyrov et al., 2004; Kemmerer et al., 2008; Boulenger et al., 2009; Shtyrov et al., 2014; Grisoni et al., 2016). Such somatotopic motor systems activation was not present in our present data on memory maintenance. However, and interestingly, there seemed to be an anterior prefrontal ‘echo’ of the previously observed word category dissociations in motor systems. This observation further strengthens the conclusion on an anterior frontal shift in verbal working memory.

### 4.1 Working memory effects

Manipulating the load of verbal working memory, the present results indicate that a distributed set of regions are important for memory encoding and maintenance. This network included regions well known to contribute to verbal working memory, especially left inferior-prefrontal/premotor and superior-temporal cortex, the cortical areas underpinning the articulatory and acoustic subparts of the phonological loop (Figure 2) (Sakai and Passingham, 2003; Thierry et al., 2003; Buchsbaum and D’Esposita, 2008). However, in the frontal lobe, the memory-active areas were not exhaustively described by the ‘Broca’s region’ label. This shows that, considering the data on memory for action related words, it is necessary to modify the Baddeley model, which emphasizes the role of Broca’s region in verbal working memory but not of other (pre)frontal sites. In particular, a prefrontal-premotor lateral focus and a dorsomedial focus of activity was characteristic of the high vs. low memory load contrast. The activation of these relatively anterior areas fits the anterior shift prediction of the neurocomputational model of the dynamics of action perception circuits.

As documented by a significant interaction of time interval by ROI, the topography of brain activation was modulated during the memory experiment. The encoding interval, during which stimuli were presented, showed strongest activation in premotor/prefrontal, left temporo-occipital and bilateral parietal areas. During memory maintenance, strongest activity was seen in left precentral and prefrontal, supplementary motor, left-perisylvian superior-temporal and bilateral parietal areas. The fact that two premotor areas, prefrontal/premotor cortex and SMA, together dominate the neurometabolic memory load effect observed during the maintenance interval suggests a role of motor systems and adjacent prefrontal cortex in the maintenance of active verbal memories semantically linked to action. While peak activation was present in premotor cortex (-46, -2, 46), this activation equally involved adjacent dorsolateral prefrontal cortex (Figure 3). The strong lateral and dorsal motor system activation seen in the present study is not typical for verbal working memory, where prefrontal and parietal activations occur together with perisylvian foci (see, for example, Paulesu et al., 1993; Jha and McCarthy, 2000; Barde and Thomspson-Schill, 2002; Sakai and Passingham, 2003; Fuster, 2009). As an explanation of this pattern of activation during action word memory, it seems plausible to consider an influence of the semantics of the stimulus materials.

### 4.2 Semantic somatotopy and verbal working memory: towards neuromechanistic integration

In the present experiment, memory load effects obtained for different action-related word categories did not appear in loci where embodied motor processes for action-related words emerged in a range of previous neuroimaging experiments (Kemmerer and Gonzalez-Castillo, 2010; Carota et al., 2012; Kemmerer, 2015). Nor did they emerge at the primary motor cortex loci where TMS pulses had elicited causal effects on action word recognition which depended on the meaning of these items (Pulvermüller, 2005). Instead, such category differences appeared when additional ROIs were defined around relative maxima of the hemodynamic load activations. These additional regions (lateral: -46 -2 44, -48 -4 50; dorsal: -4 2 58, -34 0 60) were 13-30mm anterior to where previous studies using similar sets of stimuli (for an overview, see Kemmerer and Gonzalez-Castillo, 2010; Carota et al., 2012) found differences in motor systems activations to words typically used to refer to arm and leg driven actions (lateral: -38 -20 48, dorsal: -20 -30 64, Hauk et al., 2004). At these loci, a degree of category specificity emerged in the form of a significant interaction of the factors word type and region (dorsal vs. lateral), indicating that the working memory load effect is more pronounced for leg-related action words at these dorsal premotor sites than for arm-related words. This effect was significant for dorsal regions only. The lack of a similar significant effect in lateral sites could possibly be related to the task requirements. Although care was taken to reduce the amount of motor activation in the dominant left hemisphere, the occasional button presses as well as the inhibition of button presses in non-response trials may have obscured effects in the lateral region of interest, which is close to the hand motor representation. In summary, the memory task seemed to elicit significantly more anterior frontal activation compared with reading and listening tasks and the trace of category specificity in the memory load effects appeared at a somewhat ‘disembodied’ location, with substantial anterior shift.

Recent neurocomputational modelling work may provide an explanation for our present findings. An anterior shift of activation together with reduced topographic specificity are in line with predictions of a neurocomputational model of action perception circuits, APCs, carrying word comprehension and verbal memory processes (Garagnani and Pulvermüller, 2013; Pulvermüller and Garagnani, 2014). In this model, the momentary full ignition of an APC corresponds to the recognition and semantic understanding of a single word, whereas the subsequent reverberant activity of the circuit is the material basis of verbal working memory. Although the same action perception circuit which connects knowledge about the word form with that about its meaning, including grounding information about possible referents, is active in both comprehension and memory maintenance, different parts of the circuit are respectively active. Whereas all circuit parts partake in the ignition process, only the most strongly connected circuit parts remain active after ignition and contribute to the subsequent process of activity reverberation, which implements memory maintenance. Focusing on those network parts in frontal cortex, this leads to an anterior shift of the center of gravity of activity from motor to prefrontal cortices (Figure 1). Our present results fully support such anterior shift, which was revealed by both the general memory load contrast as well as the observed traces of word category specific activations. The neurobiological explanation for such retreat of activity to prefrontal multimodal areas lies in the connectivity structure of the underlying neuroanatomical substrate (connector hub vs. less central status in the network).

In the discussion about embodied cognition and the role of disembodied processes relying on multimodal areas, the present results offer an integrated perspective based on neurobiologically realistic APCs. These circuits emerge from correlated neuronal firing related to perceptions and bodily actions. Still, they encompass modality referential sensory and motor areas as well as multimodal connector hubs. They provide a mechanism both for the ‘embodied’ grounding of word forms in the real-world entities these symbols are used to speak about and for the ‘disembodied’ retreat of memory related neuronal activity to multimodal connector hub areas. As correlations in sensory and motor information are the major driving force in the formation of APCs, this account is largely consistent with principle ideas governing the embodied cognition framework. It is important to note this, as a simplistic version of an embodied cognition approach – which would postulate the same mechanisms to be equally relevant for language understanding and verbal memory, would clearly be falsified by the present data (for discussion of oversimplified embodied positions, see, for example, Barsalou, 2016). On the other hand, a pure disembodiment perspective situating semantics exclusively in multimodal areas does not even begin to offer an explanation for the present and related results.

The mechanism underpinning the anterior cognitive shift seen in the present study, therefore, may be one of disembodiment by which the activation of a distributed action-perception circuit dynamically moves from its sensorimotor periphery and focuses on its core parts in areas that link together action and perception systems, especially in PFC. Such neurofunctional disembodiment still allows action-perception circuits to mediate between memories, actions and perceptions, so that concurrent motor movements and their causal influence on working memory (Shebani and Pulvermüller, 2013; 2018) are also compatible with, and strongly predicted by, this model. PFC may therefore be a key component in this dynamic progression of memory activity because of its strong corticocortical connectivity to both action and perception systems of the brain. Support for our interpretation also comes from recent neuroimaging evidence suggesting that the PFC is a key region for the representation of fine-grained semantic similarities among words, across categories of action-related verbs and nouns (Carota et al., 2017). Further integrating mechanisms of embodiment and disembodiment in perception, comprehension and working memory will be a challenging and exciting topic for future research into brain-grounded cognition.

## 5. Summary and conclusion

Our results show that verbal working memory for action-related words involves premotor areas, providing support for grounded and embodied theories of action-perception circuits in language and conceptual processing (Barsalou, 1999; 2008; Pulvermüller and Fadiga, 2010; Glenberg and Gallese, 2012; Kiefer and Pulvermüller, 2012; Pulvermüller, 2013). On the other hand, disembodiment was also visible in the brain responses, especially during the working memory maintenance interval, where activation was seen not at the somatotopic central loci where previous studies had found action word elicited activity, but rather in areas anteriolateral and dorsomedial to these regions, with only a trace of semantic somatotopy. This is evidence for an anterior cognitive shift in frontal cortex, possibly indicating progression from recognition and comprehension related ignition processes to reverberant memory activation within structured action-perception circuits. Even though the formation of these circuits appears to be driven by sensorimotor information, their core parts may lie in connector hubs or convergence zones of higher association cortex, including PFC, so that, during memory intervals, reverberation may gradually focus on these core parts. The results provide an important lead for neurocomputational studies that integrate memory disembodiment with current neuromechanistic theories rooted in action and perception.

## Acknowledgments

We thank Clare Cook, Rhodri Cusack, Max Garagnani, Daniel Mitchell and Yury Shtyrov for their help at different stages of this work. This work was supported by the Medical Research Council (UK) (FP, U1055.04.003.00001.01, MC_US_A060_0034, Neural basis of words), the German Research Foundation (FP, DFG grant Pu 97/22-1 on Neurosemantics), the Wellcome Trust (JBR, 103838) and the United Arab Emirates University (ZS, G00002203).

